# Survival in macrophages induces an enhanced virulence in *Cryptococcus*

**DOI:** 10.1101/2023.09.03.556094

**Authors:** Jacquelyn Nielson, Andrew Jezewski, Melanie Wellington, J. Muse Davis

## Abstract

*Cryptococcus* is a ubiquitous environmental fungus and frequent colonizer of human lungs. Colonization can lead to diverse outcomes, from clearance to long-term colonization, and to life-threatening meningoencephalitis. Regardless of the outcome, the process starts with an encounter with phagocytes. Using the zebrafish model of this infection, we have noted that cryptococcal cells first spend time inside macrophages before they become capable of pathogenic replication and dissemination. What ‘licensing’ process takes place during this initial encounter, and how are licensed cryptococcal cells different? To address this, we isolated cryptococcal cells after phagocytosis by cultured macrophages and found these macrophage-experienced cells to be markedly more virulent in both zebrafish and mouse models. Despite producing a thick polysaccharide capsule, they were still subject to phagocytosis by macrophages in the zebrafish. Analysis of antigenic cell wall components in these licensed cells demonstrated that components of mannose and chitin are more available to staining than they are in culture-grown cells or cells with capsule production induced in vitro. *Cryptococcus* is capable of exiting or transferring between macrophages in vitro, raising the likelihood that this fungus alternates between intracellular and extracellular life during growth in the lungs. Our results raise the possibility that intracellular life has its advantages over time, and phagocytosis-induced alteration in mannose and chitin exposure is one way that makes subsequent rounds of phagocytosis more beneficial to the fungus.

**IMPORTANCE:** Cryptococcosis begins in the lungs and can ultimately travel through the bloodstream to cause devastating infection in the central nervous system. In the zebrafish model, small amounts of cryptococcus inoculated into the bloodstream are initially phagocytosed, and become far more capable of dissemination after they exit macrophages. Similarly, survival in the mouse lung produces cryptococcal cell types with enhanced dissemination. In this study we have evaluated how phagocytosis changes the properties of *Cryptococcus* during pathogenesis. Macrophage experienced cells (MECs) become ‘licensed’ for enhanced virulence. They out-disseminate culture grown cells in the fish and out-compete non-MECs in the mouse lung. Analysis of their cell surface demonstrates that MECs have increased availability of cell wall components mannose and chitin—substances involved in provoking phagocytosis. These findings suggest how *Cryptococcus* might tune its cell surface to induce but survive repeated phagocytosis during early pathogenesis in the lung.

## INTRODUCTION

*Cryptococcus* (Cc) is both a ubiquitous colonizer of human lungs and the most common cause of fungal meningitis. The greatest burden of severe disease is among the immunocompromised, especially those infected with HIV (1). CNS infection can be difficult to detect before it is severe and hard to treat. A better understanding of the early stages of pathogenesis is needed in order to improve detection and prophylactic strategies.

Cryptococcal pathogenesis is thought to begin when spores or small/desiccated yeast are inhaled into the lower reaches of the lungs (2). Here they encounter alveolar macrophages, which can sometimes clear the infection. Otherwise, Cc may take up long-term, asymptomatic residence in the lung, presumably inside granulomas. If Cc is not cleared, it appears that the extent to which the host can mount a granulomatous response (via adaptive immune function) dictates the host’s ability to control fungal growth (3). Therefore, a susceptible host may develop fulminant disease either upon initial infection or after loss of control over a latent one. In either scenario, Cc growth in the presence of macrophages is required.

In the mouse model of cryptococcosis, a large inoculum of Cc delivered intravenously is capable of causing rapid CNS infection, with yeast cells entering the brain directly from the bloodstream (4-6). When mice are infected via the airway (as it is thought to occur in humans), Cc takes hold and grows in the lungs for several days prior to introducing disseminating particles (either bare yeast, yeast inside macrophages, or both) into the blood (7). These disseminating particles are far more efficient at brain dissemination than cultured Cc, since an almost undetectable blood burden (8) seeds the CNS effectively (7). Thus, time spent in the lung produces a Cc phenotype better suited to CNS dissemination than that of culture grown Cc. Similarly, in the zebrafish model we have observed that after IV inoculation, yeast cells are universally phagocytosed and spend time inside macrophages before returning to the bloodstream with enhanced capacity for virulence (9). We therefore hypothesize that phagocytosis and intracellular survival are not just barriers to cryptococcal pathogenesis, but necessary steps that enhance the ability of Cc to disseminate in the blood.

Cc encounters multiple macrophage types during its stay in the lung. Initial phagocytosis is most likely by alveolar macrophages, but other resident pulmonary macrophages, and recruited monocyte-derived macrophages are also present. Based on transcription profiling, alveolar macrophages remain a single subtype, while resident interstitial subsets fall into two types, CD14^+^ and CD14^-^ (10). Of note, there is evidence that alveolar macrophages are more likely to kill Cc than either of the interstitial types (11). The fate of Cc in macrophages also depends on activation and polarization. Most of the MAMP-receptor interactions listed above tend to induce TH1 and/or TH17 responses, which are host protective against fungi, there is evidence that Dectin-2 can induce an anti-inflammatory M2 phenotype in macrophages (12). Multiple elements of the cryptococcal cell wall can serve as microbe-associated molecular patterns (MAMPs), molecular triggers of phagocytosis which can induce both pro- and anti-inflammatory responses depending on their context and receptors. These MAMPs include chitin, β-glucan and mannose, among others. Most prominent of these is β-glucan, which is detected by macrophages via Dectin-1, complement receptor 3 (CR3) and ephrin A2 (EphA2) and tends to induce inflammation (13). In *Candida*, β-glucan is thought to be”masked” from these receptors by display of mannose, and strains unable to do so have decreased virulence (14, 15). *Cryptococcus* also expresses mannose at the cell wall where it could have a similar masking role (16). On the other hand, mannose itself is detected by phagocytes by way of Dectin-2, DC-SIGN, mannose receptor and others (17), which can induce both inflammatory and anti-inflammatory responses. All of these cell wall substances are thought to be”masked” by the polysaccharide capsule, which impedes phagocytosis by multiple mechanisms (18).

## RESULTS

### Macrophage-experienced Cryptococcus cells exhibit enhanced replication in the zebrafish larva and the mouse

In prior experiments using the zebrafish model of cryptococcosis, we observed that Cc inoculated into the bloodstream were completely phagocytosed by macrophages and neutrophils within ∼2 hours (9). After a variable amount of time (1 to 3 days), surviving yeast cells could be seen circulating in the bloodstream in small numbers. This”secondary fungemia” (since initial inoculation produced a primary fungemia) set the stage for dissemination to specific organs (including the brain) and sometimes overwhelming fungemia. The cells of secondary fungemia had essentially become more virulent. We hypothesized that time inside phagocytes had altered the virulence potential of these cells, making them capable of new interactions with the host, including dissemination and enhanced growth. To test this, we generated macrophage-experienced Cn (MECs) in vitro. We co-cultured murine J774 cells with an EGFP expressing KN99 strain (JMD163) (19) for three hours to allow adhesion and phagocytosis, then washed off any non-adherent Cn. The infected J774 cells were then incubated overnight and finally lysed so that the Cc cells could be isolated. These MECs were then inoculated into the vasculature of zebrafish larvae at our usual inoculum of 30-70 fluorescent cells per larva. MECs showed more rapid and extensive replication in the larva than either our usual YPD-cultured Cc or tissue culture-conditioned cells (control yeast treated exactly as MECs except with no J774 cells present) (Figure 1A). Host mortality was also increased, as survival of MEC-infected larvae dropped off at 4-5dpi (Fig. 1A and data not shown). To determine if this was an artifact of the J774 cell line, we repeated this experiment using human THP-1 macrophage-like cells, with same result (Fig. S1).

**Figure 1.**
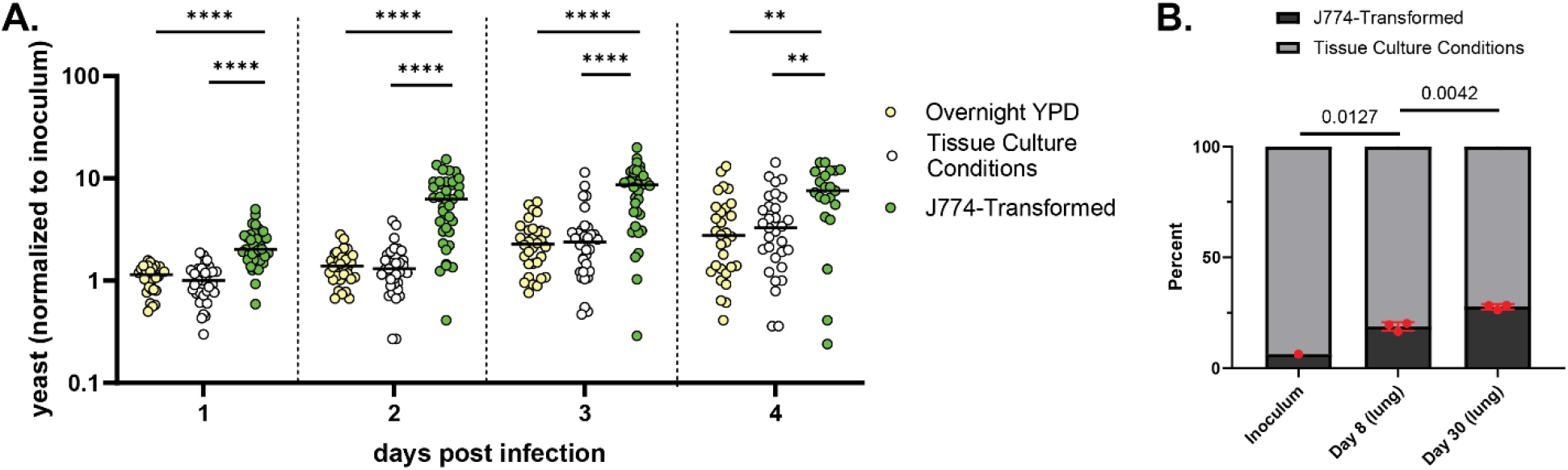
MECs show enhanced virulence in zebrafish larvae and mouse lungs. **A**. Larvae inoculated with 20-70 fluorescent yeast prepared in one of three ways. Counts were performed via live microscopy at 2hpi and 1 through 4 days post infection (dpi). Charted are daily yeast counts normalized to counts at 2hpi. Statistical comparisons represent results of Mann-Whitney tests of log transformed ratios. **B**. Comparative fungal burden of TC-conditioned (grey) and J774-Transformed Cc in the lungs of mice infected intranasally with 1x10^4^ total CFU. Statistical comparisons represent unpaired t tests of log transformed ratios.

To determine if pre-phagocytosis had the same effect in a mammalian infection, we performed a competition assay using EGFP-expressing MECs and TC-conditioned Cc expressing mRuby3 (20, 21) (JMD225, produced from the same parental strain as JMD163). These were inoculated together into A/J mice via the intranasal route. A total mixed inoculum of 1x10^4^ CFU was instilled and lungs were harvested at days 8 and 30 for fungal burden measurement. At each timepoint the proportion of EGFP-expressing cells (MECs) increased over the initial inoculum (Fig. 1B), consistent with enhanced replication by MECs in the lung. Therefore, macrophage-experienced Cn demonstrated enhanced replication inside both zebrafish and murine hosts.

### MECs have altered cell diameter and capsule production

We reasoned that the change in virulence seen in MECs would involve the most conspicuous cryptococcal virulence factor, the polysaccharide capsule. We compared cell diameter and capsule thickness of YPD grown, TC-conditioned and macrophage experienced cells using India ink staining, with results shown in Fig. 2A and B. While the cell bodies of YPD-grown yeast were modestly smaller than in the other two conditions, there were wide variations in capsule production in tissue culture conditioned and macrophage experienced yeast. Not unsurprisingly, MECs made a very thick capsule.

**Figure 2.**
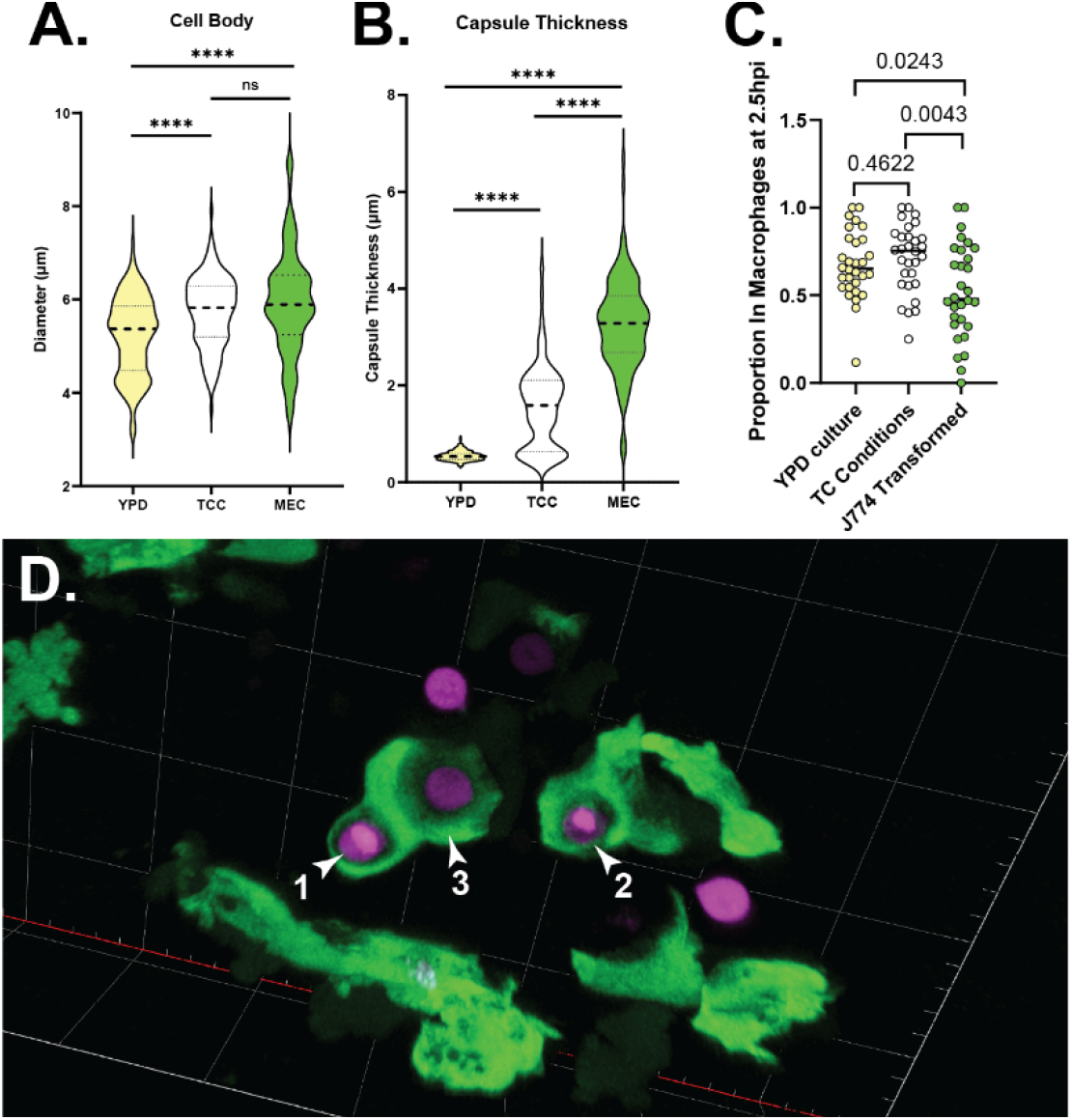
Tissue culture conditions and macrophage exposure induce increased cell body size and capsule production, with differing effects on phagocytosis. **A**. Cell body diameters for YPD-grown, TC-conditioned and J774-transformed (MEC) Cc. Diameters measured via widefield microscopy. Data from three biological replicates of at least 20 cells per condition per replicate. Statistical comparisons represent results of two-tailed unpaired t tests. **B**. Capsule diameters for same cells analyzed in panel A. Statistical comparisons represent results of two-tailed unpaired t tests. **C**. Proportion of fluorescent yeast cells inside EGFP^+^ cells (macrophages) at 2.5hpi. Statistical comparisons represent results of 2-tailed unpaired tests of log transformed ratios. **D**. EGFP^+^ cells (mpeg^+^ macrophages) phagocytosing J774-transformed Cc at ∼2.5 hpi. Yeast 1 and 2 are fully engulfed, while #3 is partially engulfed. Live confocal image taken with 40x, 1.1NA water objective. 3D rendering shown. Grid lines are 20µm apart. Animation of this data in Movie S1.

### MECs in the zebrafish undergo phagocytosis despite their encapsulation

It is known that capsule production is a primary virulence factor which prevents phagocytosis of Cc. It would follow that the enhanced growth of MECs in vivo could be due simply to extracellular survival and life. To test this hypothesis we infected *Tg(mpeg:egfp)* larvae with our red fluorescent JMD225 strain and fixed these larvae at 2.5 hours after infection to quantify rates of phagocytosis for YPD-grown, TC-conditioned, and macrophage experienced Cc. While MECs were phagocytosed significantly less than controls (Fig. 2C), there was still a surprising amount of phagocytosis. Even cells with relatively large capsules were found completely and partially engulfed by macrophages (Fig. 2D). (More detailed views of the scene in Fig. 2D are shown in Fig. S2 and Video S1).

### MECs have increased mannose and chitin exposure despite their thick capsule

One mechanism by which capsule is thought to prevent phagocytosis is by masking MAMPs on the cell wall, including mannose and chitin (16, 22). Surprised by the amount of phagocytosis heavily encapsulated MECs underwent in the zebrafish, we examined the relative exposure of these elements using fluorescent-labelled concanavalin A (ConA – which binds mannose) and wheat germ agglutinin (WGA – which binds oligomeric chitin). We analyzed micrographs of stained yeast grown in YPD, conditioned in TC and transformed in J774 cells, measuring relative fluorescence (Fig. 3A and B). Interestingly, exposure of both was greater in MECs than in TC conditioned cells. Furthermore, in MECs the ConA signal was at the periphery of the cell wall and not just the bud scars, as it was in the other conditions. (Fig. 3C). A similar peripheral pattern was seen with WGA staining (Fig. 3B). This finding was reminiscent of those by Denham et al (23), who reported a similar finding in a subset of small, dissemination-prone cryptococcal cells isolated after several days in the mouse lung. The authors proposed that extra exposure of ConA was due to the relatively thin capsules of this subset. MECs, in contrast, have variable but overall rather thick capsules (Fig. 2B). If capsule were the primary means of masking these substances, we would expect staining intensity to vary inversely with capsule thickness. This turned out not to be the case (Supplemental figure S3 A and B). Finally, in order to determine if these findings were an artifact of our fixation and microscopy methods, we analyzed the same cryptococcal strain incubated overnight in RPMI—a reliable method for inducing capsule. Induction of capsule was as expected in RPMI (Fig. S3C), but ConA and WGA staining were markedly reduced compared to MECs (Fig. S3D and E).

**Figure 3.**
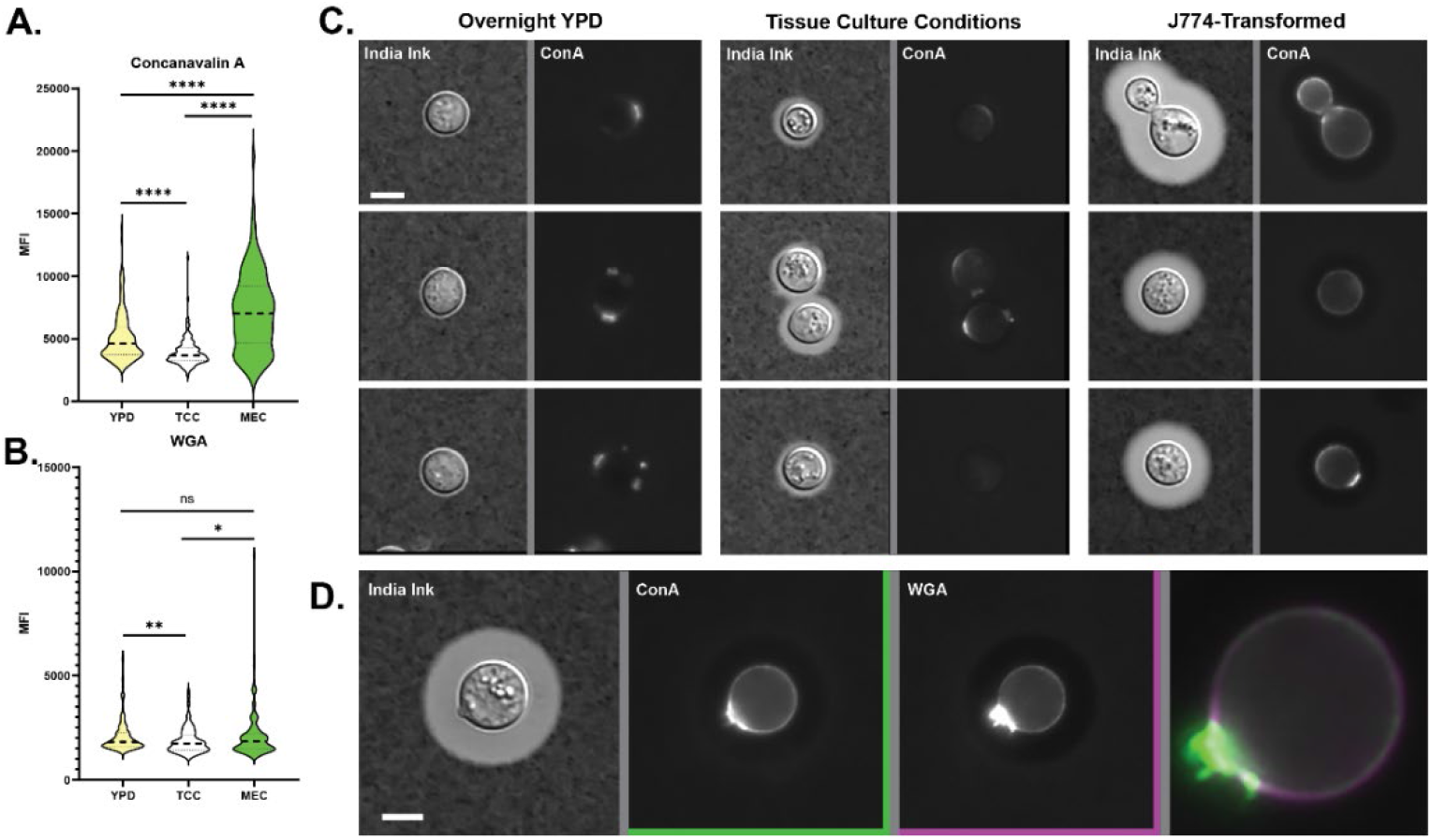
MECs have increased exposure of mannose and chitin despite production of a thick capsule. Concanavalin (**A**.) and wheat germ agglutinin (**B**.) mean fluorescence intensities for Cc cultured in YPD, TC-conditioned or J774-transformed. Statistical comparisons represent results of two-tailed, unpaired t-tests. **C**. India ink and fluorescence images of YPD-cultured, TC-conditioned and J774-transformed Cc, three examples each. Staining in the first two conditions is punctate, at bud scars, or absent, while staining of J774-transformed cells is both punctate and circumferential. ConA is tagged with Alexa Fluor 633 (Fisher #C21402). Widefield images taken with a 20x, 0.85NA objective. Exposures and intensity adjustments are uniform for all panels. **D**. Example of ConA and WGA staining overlap on a different J774-transformed cell. Note intense staining at bud scar with overlapping circumferential staining. WGA is tagged with Alexa Fluor 555 (Fisher #W32464). Widefield images taken with a 20x, 0.85NA objective.

### MECs may show enhanced brain dissemination, but TC conditioning accounts for most of this effect

In our prior investigations of brain dissemination in zebrafish, we have used cryptococcal cells grown overnight in YPD (19). In observing zebrafish infections with MECs, we noticed that brain lesions were more frequent and grew subjectively faster than in prior experiments (Fig. 4A). We have previously reported a scant 43 parenchymal lesions in a set of 77 fish (0.56 lesions per larva) (19). TC-conditioned yeast and MECs averaged more brain lesions overall (Fig. 4B), suggesting enhanced ability to enter the parenchyma. Considering our overall replication results (Fig. 1A), we were surprised to find that the brain lesion rate was indistinguishable between TC-conditioned yeast and MECs (Fig 4B). To quantify brain dissemination in greater detail, we infected *Tg6*(*kdrl:mCherry*) zebrafish (in which endothelial cytoplasm is red fluorescent) with TC conditioned or macrophage experienced yeast and followed their arrival and fates in the brain. In all we followed 78 lesions in 44 larvae infected with TC-conditioned yeast and 77 lesions in 43 larvae infected with MECs. Although there were trends toward better brain dissemination in MECs, no statistical difference was found in terms of brain lesion clearance (Fig. 4C) or day to day replication in brain lesions (Fig. 4D and E). Thus, while macrophage experience enhances the overall virulence of *Cryptococcus*, this is due primarily to enhanced growth in vivo. While MECs disseminate to the brain better than YPD-grown cells do, this difference is essentially replicated by incubation in tissue culture conditions.

**Figure 4.**
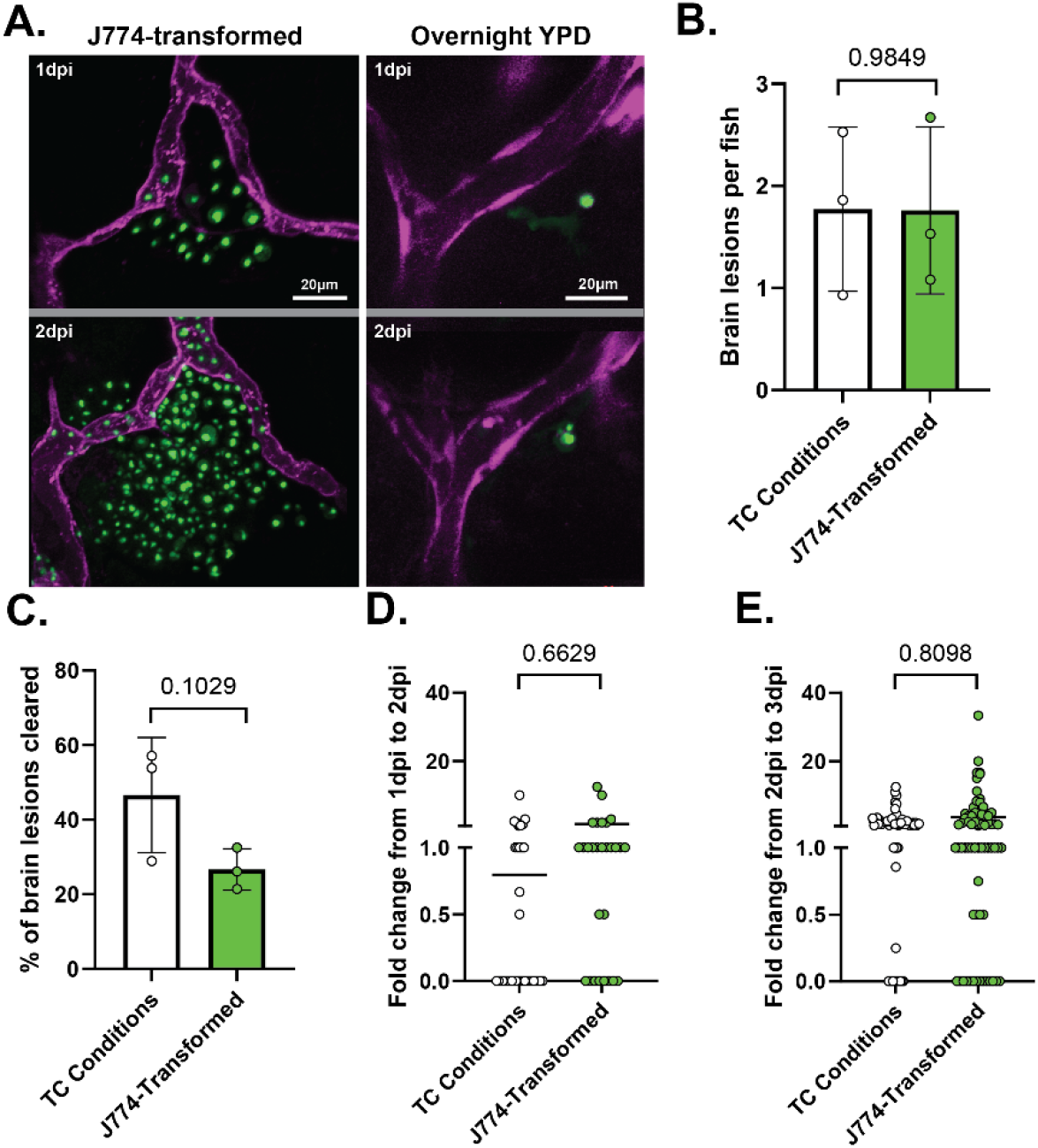
J774-transformed Cc are more effective than YPD-grown at colonizing the CNS in zebrafish, but the difference is mostly accounted for by tissue culture conditions. **A**. Yeast (green) outside the CNS vasculature (magenta) at 1 and 2dpi. Replication is greatly increased in J774-transformed Cc. Airyscan confocal image collected with 40x, 1.1NA water objective. **B**. Brain lesions per fish are no different between TC-conditioned and J774-transformed Cc. Statistical comparison for this and all other panels of this figure represent results of Mann-Whitney tests of log transformed ratios. **C**. Percent of brain lesions cleared between 1 and 3dpi. Trend is toward more clearance of TC-conditioned cells, but not statistically significant. Fold change of number of yeast per brain lesion in lesions not cleared from 1 to 2dpi (**D**.) and 2 to 3dpi (**E**.).

## DISCUSSION

The Cc-macrophage interaction is a key step in cryptococcal pathogenesis, and here we show that in vitro experience in a macrophage enhances the yeast’s ability to survive and proliferate in both the zebrafish and murine models. Evasion of host immunity is a central theme of Cc pathogenesis (24, 25), but initial survival in macrophages has implications for subsequent survival and virulence. Our data suggest that the initial interaction between yeast and macrophage licenses the yeast cells to adopt a hypervirulent phenotype. This licensed phenotype includes increased cell body and capsule size, but also a particular alteration in MAMP exposure. Interestingly, licensed cells are heavily encapsulated but still subject to significant amounts of phagocytosis. In the zebrafish we have previously reported yeast escaping from macrophages in early pathogenesis, only to be phagocytosed again (9). In the mouse lung, Cc spends 1 to 2 weeks before it is capable of bloodborne dissemination. Knowing that Cc is capable of lytic and non-lytic exit from macrophages and even direct transfer in between, it would not be surprising to find that individual yeast cells are phagocytosed and released multiple times during this period. Cc could be using selective exposure of MAMPs to sustain a sequence of alternating intra- and extracellular growth periods in the lung.

How might such a sequence benefit cryptococcal growth? Two possibilities are: a) by taking advantage of more permissive macrophage types which arrive later, and b) altering the cell surface to manipulate the nature of subsequent intracellular visits. As noted above, alveolar macrophages, likely the first encountered in the lung by Cc, are shown to be more fungicidal than the interstitial macrophages which might perform subsequent phagocytosis. Surviving and escaping alveolar macrophages might also provide Cc the opportunity to take up residence in a more permissive cell. Such a scheme for reaching permissive macrophages has been shown in mycobacterial infection (26). As for manipulating the outcome of subsequent phagocytosis, specific host receptors which initiate recognition and phagocytosis of a microbe have distinct implications for the fate of the ingested organism and the inflammatory state produced. DC-SIGN for example, one receptor which binds mannose, has a mixed reputation for inducing pro- or anti-inflammatory signaling after phagocytosis (27-29), in particular leading to expression of IL-10, an inflammation-dampening signal long associated with Cc pathogenesis. Also as noted above Dectin-2, another receptor that recognizes mannose, can induce T_H_2 immunity in certain circumstances (12). Therefore by increasing mannose exposure, initial phagocytosis could well be inducing changes which alter the outcome of subsequent macrophage encounters.

A recent mouse study has revealed that average cryptococcal cell size gradually decreases in the 10 to 14 days between lung infection and bloodborne dissemination (23). The authors identified in the lung a distinct population of the smallest Cc cells most capable of dissemination to the brain. What process drives this differentiation? Our data suggest that a single sojourn inside of a macrophage is not enough to account for it. We show that MECs, with their single night of macrophage experience, are better at brain dissemination than YPD-grown Cc, but their advantage over TC-conditioned cells is minimal at best. The best characterization of the MECc virulence advantage is sheer in vivo replication, whether in the bloodstream and brain of the fish or in the lungs of the mouse.

Phagocytosis is a vital step in cryptococcal pathogenesis. Any yeast which ultimately reach the CNS to cause severe disease have spent time inside a macrophage, or are daughters of cells which have. While the ability to survive within, and escape macrophages is key to the virulence potential of this pathogen, our results raise the possibility that macrophages are more than an opposition to pathogenesis, but a necessary collaborator.

## MATERIALS AND METHODS

### Zebrafish care and maintenance

Adult zebrafish were kept under a light/dark cycle of 14 h and 10 h, respectively. Larval zebrafish were incubated at 28.5°C in E3 buffer (30), switching larvae to E3-MB containing 0.2 nM 1-phenyl-2-thiourea (PTU) (Sigma-Aldrich) at 18-24 hours post fertilization to inhibit pigment formation. All larvae were manually dechorionated between 24 and 30 hpf. Prior to microinjection or imaging, larvae were anesthetized in E3-MB containing 0.2 mg/ml Tricaine (ethyl 3-aminobenzoate; Sigma-Aldrich). For prolonged time lapse imaging, larvae were mounted in 1% low-melting point agarose, 0.2 mg/ml Tricaine, and 0.2nM PTU (final concentrations) on a coverglass-bottom dish. All adult and larval zebrafish procedures were in full compliance with NIH guidelines and approved by the University of Iowa Institutional Animal Care and Use Committee (Protocol #0102075-002).

### Transgenic Zebrafish Lines

*Tg(kdrl:mCherry-CAAX)* (31) is designated y171Tg at the Zebrafish International Resource Center (ZFIN) and was kindly provided by Daniel Castranova. *Tg(mpeg:egfp)* (32) is designated gl22Tg and was kindly provided by Dr. Anna Huttenlocher. Transgenic lines are maintained by outcrossing to AB wildtype fish every other generation. ABs are obtained from ZFIN regularly to maintain hybrid vigor.

### Cryptococcal strains and growth conditions

KN99α was kindly provided by Dr. Kirsten Nielsen. Cultures were handled using standard techniques and media as described previously (33, 34).

### Construction of fluorescent *Cryptococcus neoformans* strains

A nuclear localized mRuby3 expression strain (JMD225) was constructed using CRISPR, similar to the prior construction of the green fluorescent JMD163 strain (19). mRuby3 was chosen based on its impressive fluorescence in Spencer et al (21). The mRuby3 construct was amplified from pGWKS7 (AddGene #139414) using primers JMD228 (5’ – cat cta tca cAT GGT CTC CAA GGG CGA G – 3’) and JMD231 (5’ – tgg cgg atc cCT TGT AGA GCT CGT CCA TGC – 3’) (Lower case letters indicate overlap region for fusion PCR). An 80 amino acid nuclear localization sequence was amplified from pCH1227 (9) using primers JMD233 (5’ – gct cta caa gGG ATC CGC CAA GAA GGA TG – 3’) and JMD207 (5’ – agt gac gga tta gTA TCA TCA CGC CAC ACC C – 3’). A histone 3 promoter was amplified from pCH1227 using primers JMD212 (5’ – tca acg att ttg GAG CTC GGC AGA TAC GAT ATG – 3’) and JMD234 (5’ – aac gat ttt gGT GAT AGA TGT GTT GTG GTG – 3’). pH3mchSH2 (AddGene #101053, (Upadhya, Lam et al. 2017) was used as a source for the SH2 flanking regions and nourseothricin resistance cassette. The 5’ flanking region was amplified using primers JMD175 (5’ – GAT GTC TTA AAT AAA AGT TTG GAG TCT AGA GCT GGT CTT TAT CAT G – 3’) and JMD205 (5’ – atc tgc cga gct cCA AAA TCG TTG AAC CCG C – 3’). The 3’ flanking region and nourseothricin resistance cassette were amplified using primers JMD176 (5’ – CCT TTG AAT TAC ATG GAG TAG TTG CAA GGC CC – 3’) and JMD206 (5’ – ggc gtg atg ata CTA ATC CGT CAC TCC TTT TTT C – 3’). PCR products were then joined using the NEBuilder HiFi DNA Assembly Kit and electroporated into KN99α as detailed in (35). Colonies expressing red fluorescence were collected and genomic DNA isolations were performed as per (36). The samples were then screened by PCR for proper insertion of the construct into the SH2 locus. Primer pair JMD170 (5’ – CCT TCA TCA ACA TTC TGA CGC TTT GTT TCG TGA AGC TTG T – 3’) and JMD27 (5’ – CGT TGT TTC AGG CCT GCG GAT G – 3’) produce a 1619bp band from an intact SH2 locus or a 6272bp band from an SH2 site with the mRuby3 construct in place. Positive candidates were then compared with our GFP strain (JMD163) for growth in culture and the strain with similar growth (JMD225) was used in the experiments here.

### *Cryptococcus* macrophage transformation

#### J774A

The mouse macrophage cell line J774A.1 (ATCC TIB-67) was kindly provided by Dr. Melanie Wellington. A confluent T-75 flask of cells exhibiting normal morphology and cell adhesion was split 1:2 in 15 mL DMEM supplemented with 10% FBS, 1% Glutamax, and 1% penicillin streptomycin and allowed to grow overnight. *Cryptococcus* strains were cultured in 50 mL liquid YPD overnight. The *Cryptococcus* cells were washed three times in sterile DPBS. 2x10^9^ *Cryptococcus* cells/mL were opsonized with the 18B7 monoclonal antibody (provided by Dr. Damian Krysan) for 30 minutes prior to co-incubation with macrophages. J774A.1 medium was replaced with 15 mL fresh medium and 5x10^8^ *Cryptococcus* cells (MOI 20), followed by a 3 hour incubation at 37°C with 5% CO_2_. Simultaneously, 5x10^8^ *Cryptococcus* cells were added to 15 mL J774A.1 culture medium in a T-75 culture flask and incubated at 37°C with 5% CO_2_ as a control. The macrophage/*Cryptococcus* co-culture was then washed 3 times with warm DPBS to remove non-adherent yeast, then replenished with 15 mL fresh J774A.1 medium and allowed to incubate overnight at 37°C with 5% CO_2_. To lyse the macrophages, 5 mL 10% Triton X-100 in DPBS (final concentration of 2.5%) was added to the co-culture and the control flask then incubated for 20 minutes at room temperature while rocking. *Cryptococcus* cells were then centrifuged and washed twice with sterile DPBS before use. ***THP-1*** The human monocyte cell line THP-1 (ATCC TIB-202) was also kindly provided by Dr. Melanie Wellington. Cells were harvested from confluent flasks, then seeded in a T-75 flask at a density of 3x10^5^ cells/mL in 15 mL RPMI supplemented with 10% FBS, 1% Glutamax, 1% penicillin streptomycin, 10mM HEPES, and 1mM sodium pyruvate. Cells were primed with 100 ng/mL PMA and allowed to incubate for 2 days at 37°C with 5% CO_2_ (37). Media was then replaced with culture medium without PMA and cells were incubated for an additional day at 37°C with 5% CO_2_. *Cryptococcus* strains were cultured in 5 mL liquid YPD overnight. The *Cryptococcus* cells were washed three times in sterile DPBS. 1x10^8^ *Cryptococcus* cells/mL were opsonized in pooled human serum (MP Biomedicals) for 30 minutes and washed 3 times with DPBS prior to co-incubation with macrophages. THP-1 medium was replaced with 15 mL fresh medium and 9x10^6^ *Cryptococcus* cells (MOI 20), followed by a 3 hr incubation at 37°C with 5% CO_2_. Simultaneously, 9x10^6^ *Cryptococcus* cells were added to 15 mL THP-1 culture medium in a T-75 culture flask and incubated at 37°C with 5% CO_2_ as a control. The co-culture was then washed 3 times with warm DPBS to remove non-adherent *Cryptococcus* cells, then replenished with 15 mL fresh THP-1 medium and allowed to incubate overnight at 37°C with 5% CO_2_. To lyse the macrophages, 5 mL 10% Triton X-100 in DPBS (final concentration of 2.5%) was added to the co-culture and the control flask then incubated for 20 minutes at room temperature while rocking. *Cryptococcus* cells were then centrifuged and washed twice with sterile DPBS before use.

#### *Cryptococcus* Lectin Staining

Macrophage transformed, culture medium transformed, and overnight YPD cultured *Cryptococcus* cells were washed twice in sterile DPBS and resuspended in DPBS at a concentration of 1x10^8^ cells/mL. *Cryptococcus* cells were then fixed in 2% paraformaldehyde in DPBS for 12 minutes at room temperature and washed three times with sterile DPBS. Cells were stained with concanavalin A (Con A) tagged with Alexa Fluor 633 (Fisher #C21402) and wheat-germ agglutinin (WGA) tagged with Alexa Fluor 555 (Fisher #W32464) as previously described (23) in addition to calcofluor white. Briefly, cells were incubated in WGA for 30 minutes in the dark at room temperature while rotating. Following incubation, 50 ug/mL Con A was added to the cells and incubated for an additional 3 minutes. Cells were then centrifuged and washed twice in DPBS, then combined 1:1 with India ink for imaging.

#### Zebrafish infection by microinjection

Single colonies from 4°C stocks were inoculated into YPD medium and incubated shaking overnight at 30°C. Additionally, *Cryptococcus* cells were transformed in macrophages or culture medium as described above. YPD cultures were washed twice with sterile DPBS before use. All cultures were diluted in DPBS containing 10% glycerol and 2% PVP-40 (polyvinylpyrrolidine, Sigma Aldrich) (38) to an OD_600_ of 5.0 in a 1:10 dilution of phenol red. After manual dechorionation of embryos at ∼28 hpf, IV inoculations were performed as previously described (39), with the alteration that larvae were positioned on a 3% agarose plate formed with holding grooves as described in (30). Initial inoculum was documented by direct microscopic observation and only larvae with initial inoculum between ∼30 and 70 fluorescent yeast cells were used.

#### Live zebrafish microscopy

Confocal imaging was performed on a Zeiss Axio Observer.Z1/7 body equipped with a Zeiss Airyscan detector. All channels were collected using Airyscan Multiplex settings with default processing. Cryptococcal India ink and lectin staining images (Figures 3 and S3) were captured using a Zeiss LD C-Apochromat 60x/1.1 Oil Korr UV VIS IR objective. Correlative DIC images in Fig. S2 were captured shortly after the confocal collection in widefield mode using a Zeiss 40x/1.1 Water corrected Plan Apochromat objective and re-scaled manually to overlay with confocal images taken with the same objective. All other images were captured using a Zeiss Plan-Apochromat 20x/0.8 objective. Live imaging was performed with larvae anesthetized in tricaine as previously described (39) and simply resting on the bottom of a glass-bottom dish or immobilized in 1% low melt agarose. For collection of large numbers of events, a combination of widefield and confocal imaging was used to create scout and detail images for later analysis. Widefield imaging was performed using the same Zeiss optical setup, with image capture using a Hamamatsu Flash4.0 V3 sCMOS camera. Widefield fluorescence excitation was generated with a Colibri 7 type RGB-UV fluorescence light source. Filter set was Zeiss set 90 LED.

#### Fixed zebrafish microscopy

Larvae were inoculated with cryptococcal cells prepared as described elsewhere and euthanized in 250mg/L Tricaine-S at 2.5hpi. Larvae were then fixed with 4% paraformaldehyde overnight at 4°C. Prior to imaging they were washed repeatedly in 1X PBS and stored at 4°C until viewing. Images for analysis of phagocytosis were collected in widefield mode using a Zeiss Plan-Apochromat 20x/0.8 objective. Confocal image in Fig. 2D and Video S1 was collected using Airyscan multiplex settings and default Airyscan processing.

#### Image processing

Images were processed with Zeiss Zen software applying default Airyscan processing followed by z-stack alignment if needed. Single slices, orthogonal projections and 3D renderings were produced in Zen software and exported as TIF files for assembly using Adobe Photoshop and Illustrator. 3D renderings were done with default maximum settings in Zeiss Zen.

#### Murine virulence studies

*Cryptococcus* cells were transformed in J774A.1 macrophages or culture medium and collected as described above. To differentiate cells from each condition, GFP expressing *Cryptococcus* cells (JMD163) were incubated with macrophages and mRuby3 expressing *Cryptococcus* cells (JMD225) were incubated in culture medium. Following collection, both cultures were diluted in DPBS containing 10% glycerol and 2% PVP-40 (polyvinylpyrrolidine, Sigma Aldrich) (38). 8-10 weeks old A/J female mice (JAX) were inoculated with a 1:1 mix of each *Cryptococcus* strain as described in (40). Briefly, mice were inoculated with 5x10^3^ CFU of each strain via intranasal route (1x10^4^ CFU total). Each group contained 3 mice. Brains and lungs were collected 8 days post inoculation and homogenized in sterile PBS. Homogenates were diluted in a 10-fold dilution series, plated on YPD, and incubated at 30°C for 48h. Colonies were counted using a Zeiss SteREO V8 microscope with Zeiss achromat S 0.63x Reo objective and Lumencor SOLA light engine, to differentiate GFP and mRuby3 expressing cells.

## FIGURES

**Figure S1.**
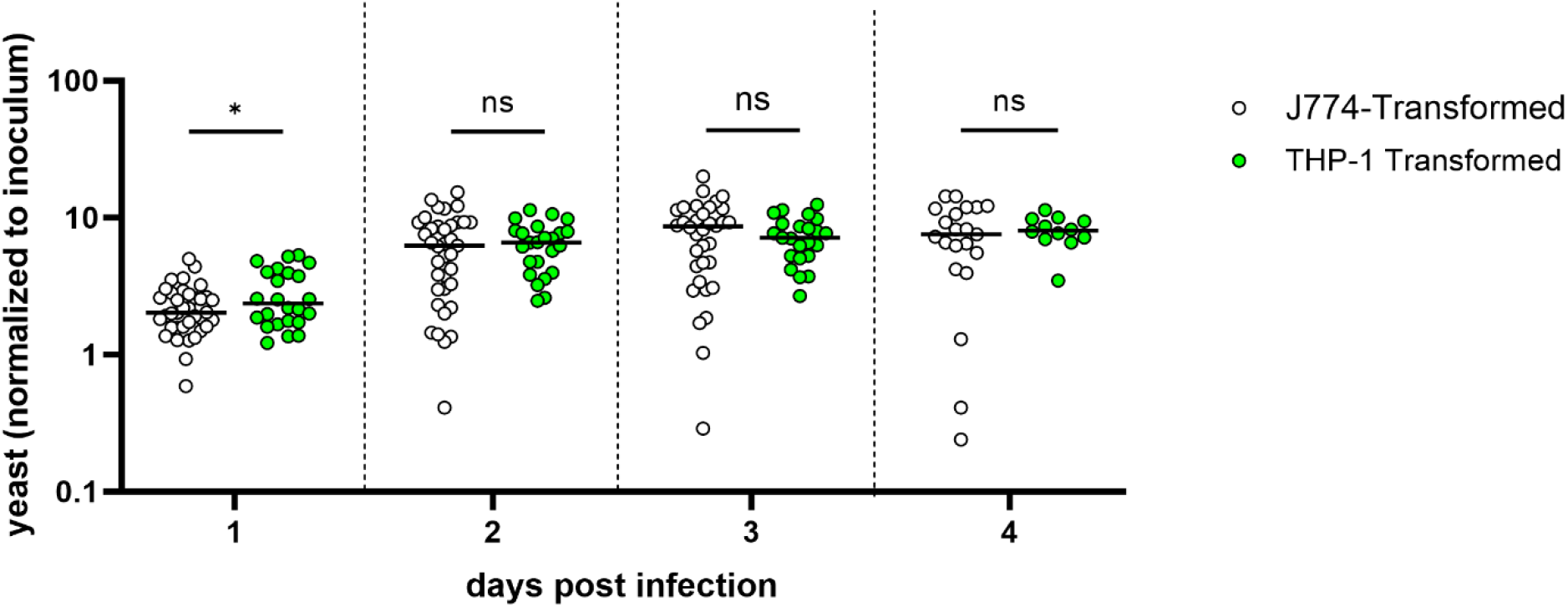
Enhanced virulence in fish is seen in MECs from both murine and human cell lines. Larvae inoculated with 20-70 fluorescent yeast each as in Fig 1A. Counts were performed via live microscopy at 2hpi and 1 through 4 days post infection (dpi). J774-transformed data is the same as in Figure 1A, compared to THP-1 data separately for clarity. Charted are daily yeast counts normalized to counts at 2hpi. Statistical comparisons represent results of Mann-Whitney tests of log transformed ratios.

**Figure S2.**
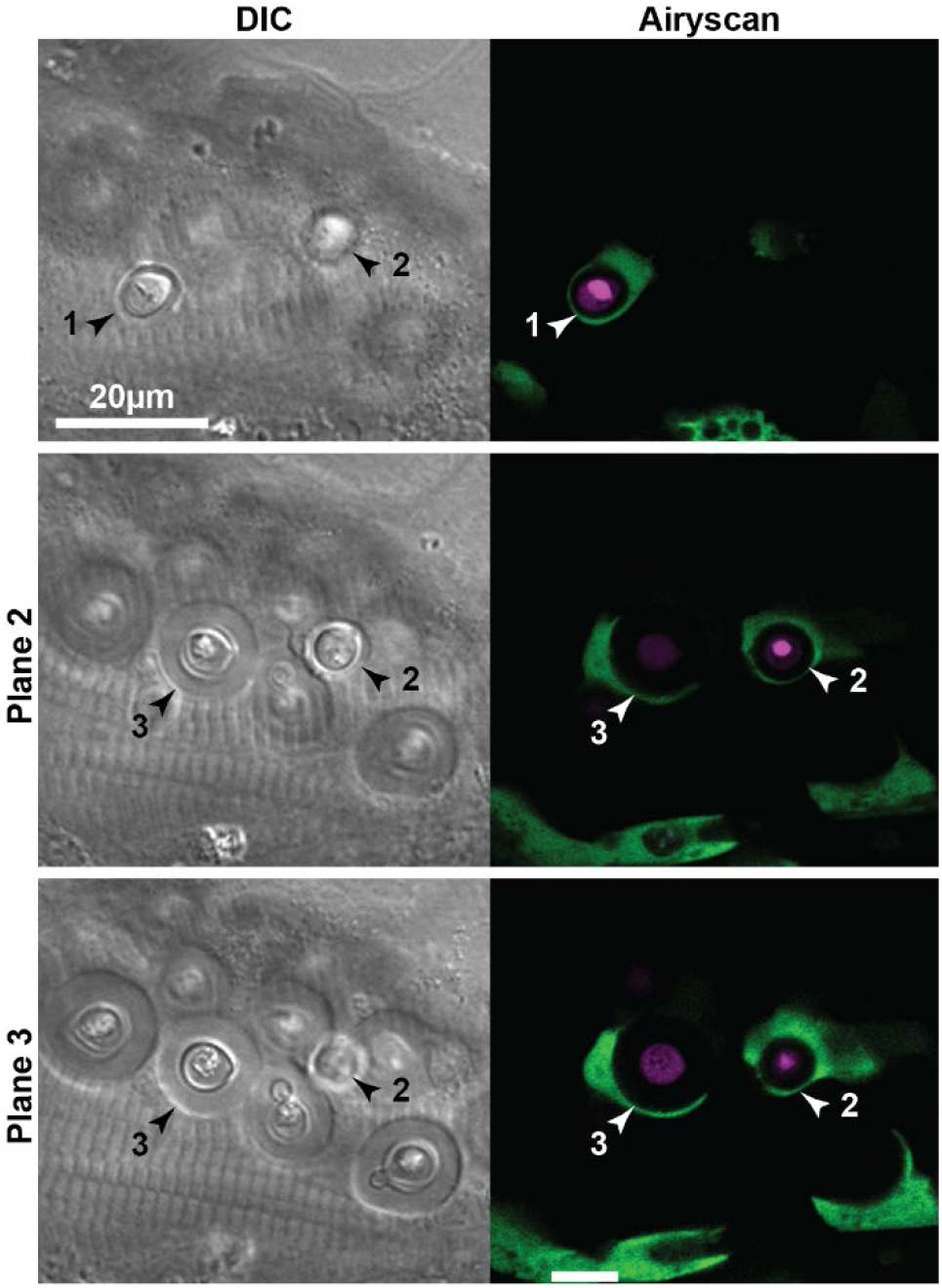
Single slice images of MEC phagocytosis with correlating DIC images. Same image set as 3D rendering in Figure 2D, with addition of matching DIC images taken within minutes of confocal ones. Individual yeast are numbered to match Figure 2D. Animation of this dataset in Movie S1.

**Figure S3.**
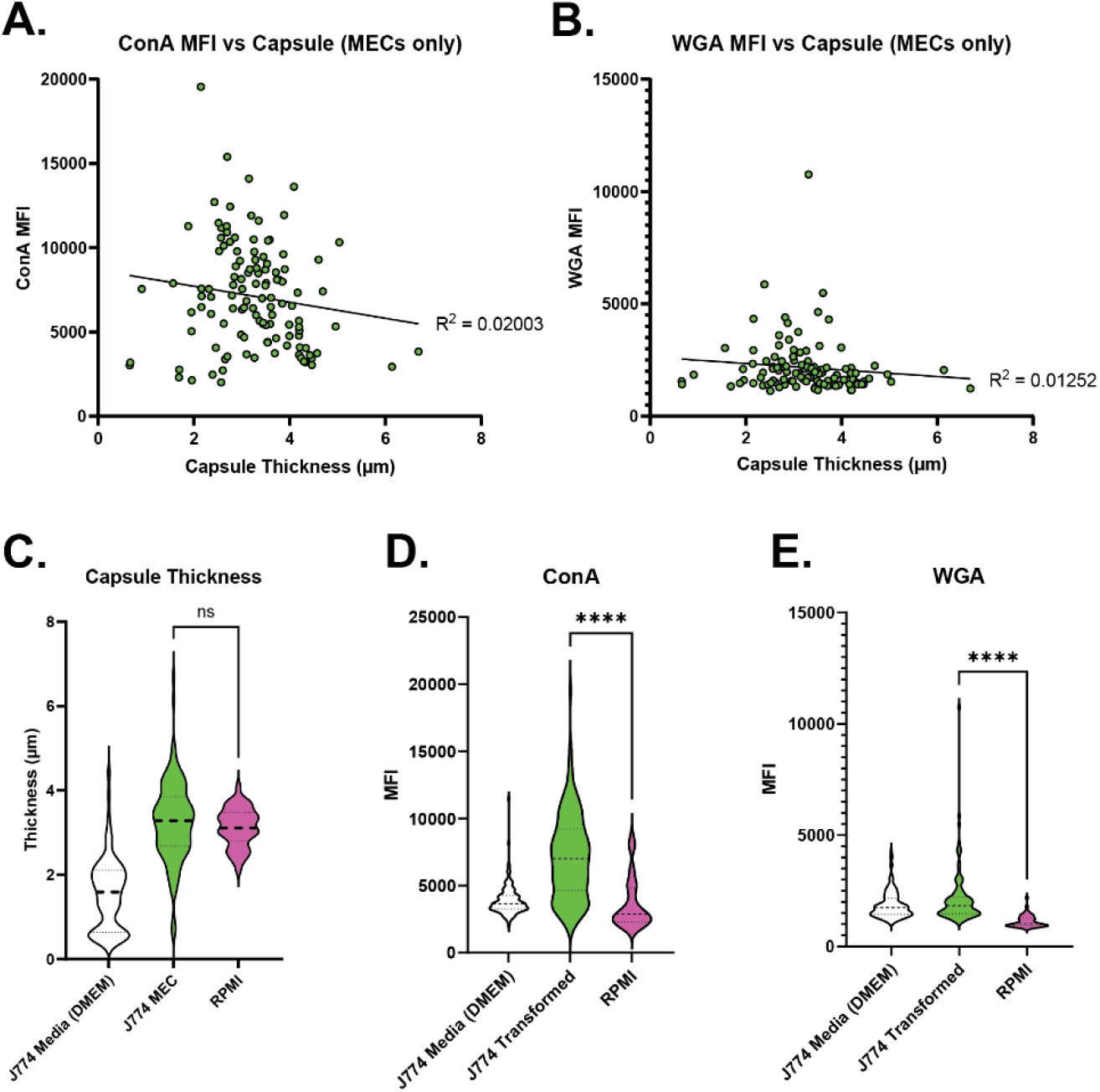
Capsule thickness and ConA, WGA exposure in MECs and comparison to RPMI-induced encapsulated Cc. **A**., **B**. No linear correlation between ConA (**A**.) and WGA (**B**.) exposure and capsule thickness. R^2^ plotted via simple linear regression. **C-E**. Capsule thickness (**C**.) is similar in J774-transformed Cc and Cc with capsule induced with RPMI, But exposure of ConA (**D**.) and WGA (**E**.) is less. J774 media and MEC data same as in Figure 2B, re-displayed in this context for clarity. Statistical comparisons represent results of two-tailed unpaired t tests.

**Movie S1**. Same image depicted in Figures 2D and S2, with rotation to emphasize full internalization of 2 of 3 yeast cells indicated in those figures.

